# Identification of Pirin as a Molecular Target of the CCG-1423/CCG- 203971 Series of Anti-Fibrotic and Anti-Metastatic Compounds

**DOI:** 10.1101/458554

**Authors:** Erika M. Lisabeth, Dylan Kahl, Indiwari Gopallawa, Sarah E. Haynes, Sean A. Misek, Phillip L. Campbell, Thomas S. Dexheimer, Dinesh Khanna, David A. Fox, Xiangshu Jin, Brent R. Martin, Scott D. Larsen, Richard R. Neubig

**Affiliations:** Department of Pharmacology & Toxicology; Department of Medicinal Chemistry; Department of Chemistry; Department of Internal Medicine, Division of Rheumatology; Department of Biochemistry, Michigan State University, East Lansing, MI, USA; Vahlteich Medicinal Chemistry Core, College of Pharmacy, University of Michigan, Ann Arbor, MI, USA

## Abstract

A series of compounds (including CCG-1423 and CCG-203971) discovered through an MRTF/SRF dependent luciferase screen has shown remarkable efficacy in a variety of *in vitro* and *in vivo* models, including melanoma metastasis and bleomycin-induced fibrosis. Although these compounds are efficacious, the molecular target is unknown. Here, we describe affinity isolation-based target identification efforts which yielded pirin, an iron-dependent co-transcription factor, as a target of this series of compounds. Using biophysical techniques including isothermal titration calorimetry and X-ray crystallography, we verify that pirin binds these compounds *in vitro*. We also show with genetic approaches that pirin modulates MRTF-dependent SRE.L Luciferase activation. Finally, using both siRNA and a previously validated pirin inhibitor, we show a role for pirin in TGF-p induced gene expression in primary dermal fibroblasts. A recently developed analog, CCG-257081, which co-crystallizes with pirin, is also effective in the prevention of bleomycin-induced dermal fibrosis.

## Introduction

Myocardin-Related Transcription Factor (MRTF) and Serum Response Factor (SRF) are transcription factors activated downstream of the Rho family of GTPases, which regulates actin cytoskeleton and motility.^1,2^ The MRTF/SRF complex activates a gene transcription program involved in the expression of structural and cytoskeletal genes, as well as pro-fibrotic genes.^3^ This mechanism feeds back on cell motility control and has been strongly implicated in cell migration and proliferation. Rho/MRTF/SRF signaling has also been linked to melanoma metastasis and to fibrotic pathological mechanisms.^4,5^ In 2007, we reported the first tool compound inhibitor of the Rho/MRTF/SRF pathway, CCG-1423.^6^ It was identified in a pathway screen using a modified MRTF-dependent serum-response element (SRE.L) luciferase assay.^6^ CCG-1423 was shown to have anti-migratory and anti-proliferative effects in prostate cancer and melanoma cells *in vitro*.^6^ It has been extensively used as an “MRTF inhibitor” in many biological contexts, including reduction of endothelial cell migration and angiogenesis^7^, and it has efficacy in multiple preclinical disease models, including improvement of glycemic control of insulin-resistant mice^8^ and reduction of mouse peritoneal fibrosis.^10^

Subsequent chemical modification and structure-activity relationship (SAR) studies based on the cell-based SRE-luciferase assay yielded the nipecotic acid derivatives CCG-100602 and CCG-203971^11–12^, which removed the labile and potentially reactive N-alkoxybenzamide functionality of CCG-1423. These analogs provided improved tolerability *in vivo* as exemplified by their capability to reduce bleomycin-induced skin and lung fibrosis^13,14^ and melanoma lung metastasis in two separate preclinical murine models.^15^ Further improvements to the series resulted in CCG-222740 and CCG-232601, which prevented ocular fibrosis and skin fibrosis *in vivo*.^16–17^ Recent optimization for metabolic stability yielded CCG-257081, which has improved pharmacokinetic properties but it has not been tested in *vivo* (Figure 1A-D).^17^

**Figure 1.**
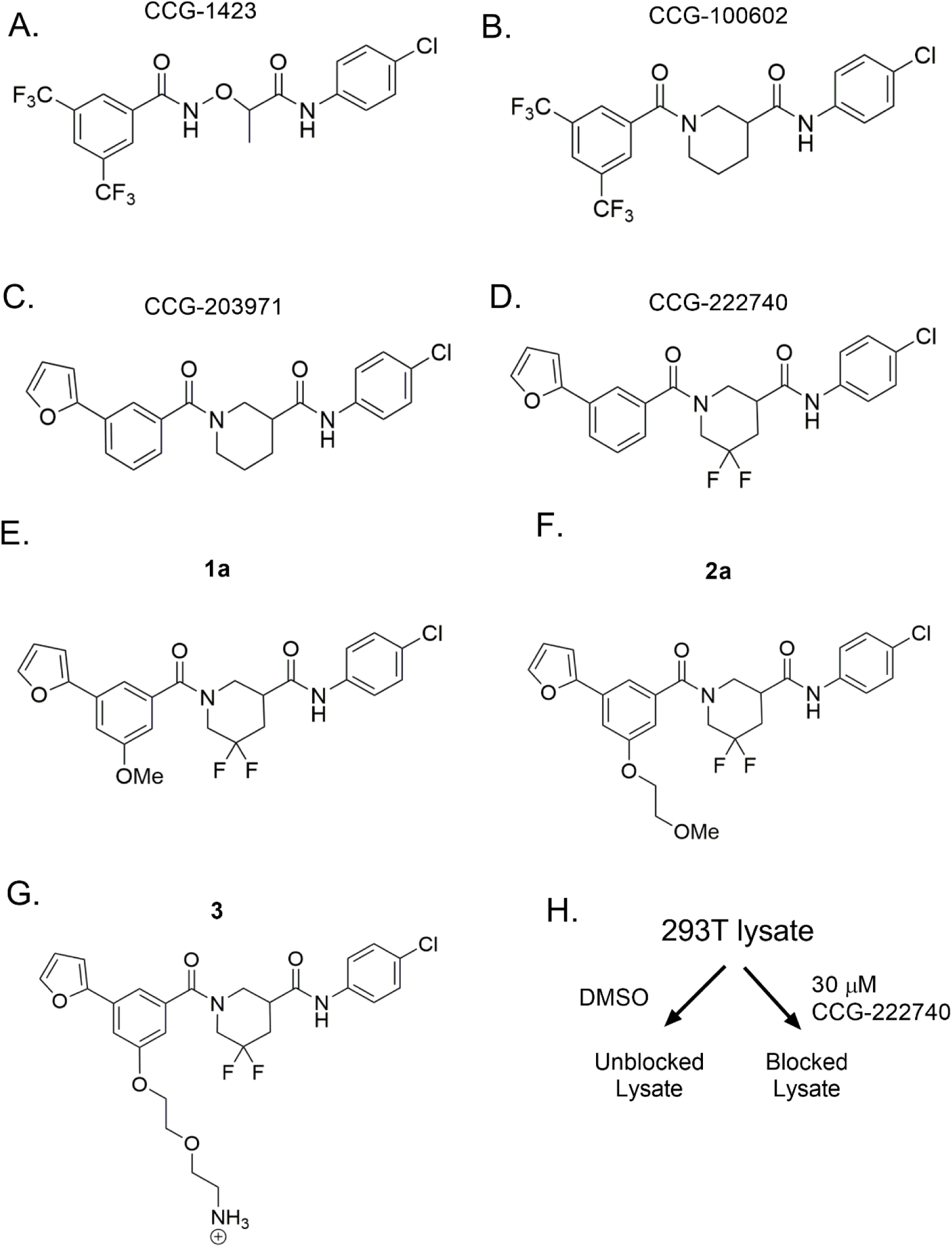
Key analogs of the Rho/MRTF/SRF-mediated gene transcription inhibitors and probe development analogs. (A-D) Important analogs within the series that have led to the development of biologically active inhibitors. (E-G) Probe mimic analogs of CCG-222740 (**1a** and **2a**) retained activity in Q231L Gα_12_ driven SRE.L Luciferase (2 μM and 2.3 μM, respectively). Based on this activity, the 5-position of the 3-furyl aromatic ring was selected for PEG-linker attachment for NHS-agarose beads (**3**). (H) HEK 293T lysates were incubated with probe **4** bound to NHS beads, with or without preblocking lysates with 30 μM of CCG-222740. Proteins bound to the beads were then analyzed by mass spectrometry.

A major limitation to the development of this series of inhibitors has been the unknown molecular mechanism. Prior target identification campaigns for this series have been inconclusive. For instance, it has been reported that CCG-1423 can interact with the N-terminal nuclear localization sequence (NLS) in the RPEL domain of MRTF-A and block MRTF-A nuclear translocation by inhibition of the interaction between MRTF-A and importin α1/β1.^18^ CCG-1423 was also shown to bind directly to MICAL-2—an atypical intranuclear actin regulatory protein that mediates SRF/MRTF-A-dependent gene transcription.^19^ MICAL-2 is proposed to function by inducing redox-dependent depolymerization of nuclear actin, ultimately decreasing nuclear G-actin and increasing MRTF-A in the nucleus.^19^ Moreover, a photolysis photoprobe was synthesized and specifically labelled an approximate 24 kDa protein band in PC3 cells; however, the identity of the protein band was never validated by mass spectrometry.^20^ Finally, a microarray analysis of gene transcription changes in PC3 cells treated with CCG-1423 had significant overlap with effects of Latrunculin B — an actin polymerization inhibitor — as well as effects on cell cycle, ER stress and metastasis gene networks, suggesting shared biological targets between these two inhibitors.^21^

Despite multiple potential molecular targets for the series, there has been no robust biophysical evidence presented. Consequently, we undertook an unbiased mass spectrometry-based target identification approach and identified pirin — a conserved iron binding co-transcription factor that has not previously been associated with Rho/MRTF signaling or fibrosis — as the top biological target candidate for the series. Pirin was the most highly enriched protein upon affinity pulldown with an immobilized active compound. It was subsequently validated through various biophysical techniques, such as X-ray crystallography and isothermal titration calorimetry (ITC), which show that our compounds bind directly to pirin. Moreover, we show that pirin is involved in MRTF/SRF dependent pro-fibrotic signaling. Finally, we also report anti-fibrotic effects of CCG-257081 in a bleomycin-induced skin fibrosis model.

## Results and Discussion

### Identification of pirin as a potential target of CCG-222740

To develop an affinity matrix for target enrichment, CGG-222740 was used as the starting chemical scaffold, mostly because of its superior inhibition of TGFp-induced ACTA2 gene expression as compared to CCG-203971.^17^ To identify optimal linker placement, we performed a methoxy scan followed by an ethoxymethoxy scan on each aryl ring (Supplemental Table I). Taking into consideration the flat SAR, retained potency, efficacy, and availability of starting material, we chose to attach the PEG linker at the 5-position of the 3-furyl phenyl ring. Since attaching either small (**1a**) or large (**2a**) functional groups at that position did not markedly reduce activity, we were confident that attaching the probe linker and resin there would allow for the probe to maintain acceptable binding affinity to the biological target(s) (Figure 1E-G). **3** was synthesized and linked to NHS-agarose beads using solid phase amine coupling chemistry followed by blocking of residual reactive groups with ethanolamine to produce **4** (Supplementary Schemes 1-3), which was subsequently used as our affinity pulldown probe.

The agarose-bound probe was mixed with whole cell lysates from HEK-293T cells containing DMSO (unblocked) or 30μM CCG-222740 (blocked)(Figure 1H). Beads were then washed several times with cold PBS (adapted from^22^). Pulldown mixtures were analyzed after trypsin digestion and subsequent proteomic analysis (see Supplemental Methods). Comparison between the normalized peptide abundance in the DMSO control lysates and the CCG-222740 treated cell lysates revealed one protein that stood out with 5-fold greater peptide abundance in the DMSO control lysates and a p-value of 1.3 e^-13^ (Table I and Supplemental Table II). Consistent with the known activity of our compounds to modulate gene transcription, it was encouraging that the enriched protein was a known co-transcription factor — pirin.

**Table I.**
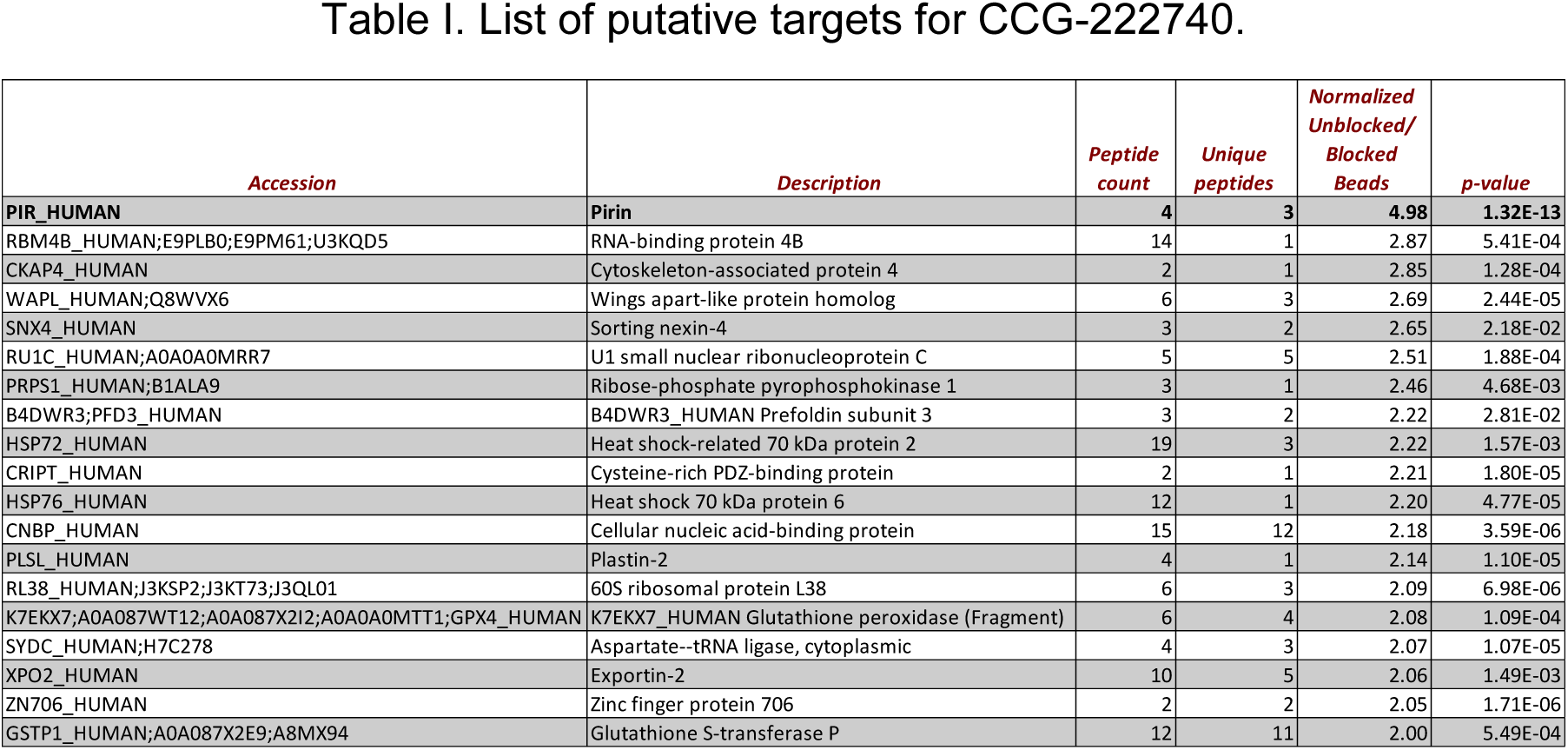
Proteins with >2 fold enrichment. HEK 293T lysates were blocked either with 30 μM CCG-222740 or DMSO control, and incubated with NHS-agarose bound CCG-262545. Peptides aligning with proteins and counts were compared in unblocked and blocked samples, and arranged in order of decrease in fold differences between the two samples. All protein samples with >2 fold differential peptide amounts are shown.

### Pirin binds to CCG-222740 and CCG-257081

Pirin is a well-conserved protein originally identified in a yeast two-hybrid screen with the transcription factor NF-1.^23^ Pirin has also been implicated in NF-κB signaling through its interaction with Bcl3, a NF-κB co-activator, and through direct interaction with a p65/p65 homodimer.^24–26^ Pirin is expressed ubiquitously, and basal expression is, in part, regulated through Nrf2-regulated gene transcription.^27^ Two drug discovery campaigns have previously been described for pirin. The first published campaign used fluorescently labeled pirin to identify small molecules in an immobilized compound library that directly bound to pirin; however, the cellular effects of the identified compound, TphA, were modest.^25^ A second study identified the compound CCT251236 while screening for inhibitors of the Heat Shock Factor 1 (HSF1) transcription pathway.^28^ Their target identification efforts yielded pirin as the target of CCT251236, which has efficacy in an ovarian cancer model *in vivo* and inhibits melanoma migration *in vitro*.^28^ The structural comparison between CCT251236 and our series of Rho/MRTF/SRF-mediated gene transcription inhibitors reveals high similarity. This supports our initial rationalization of pirin as a biological target for our series of inhibitors.

To understand whether our series of compounds and pirin regulate a similar set of genes, we took a bioinformatics approach to compare a gene expression signature of CCG-1423 target genes with 12,922 MSigDB or in-house gene sets (Supplemental Table V). Only 347 (2.7%) of the gene sets showed a statistically significant correlation with the CCG-1423 regulated genes at the Bonferroni-corrected p-value (p < 3.87×10^-6^). A siPirin gene set was the most significantly enriched (p = 4.44×10^-72^), with an overlap of 44 genes with the CCG-1423 signature (Figure 2A and Supplemental Table IV). By comparison, the overlap between our in-house MRTF signature and the CCG-1423 gene signature was 8 genes (data not shown). Therefore, the bioinformatics analysis, comparing the CCG-1423 microarray dataset and the siPirin microarray dataset, was very encouraging and consistent with the idea that pirin may be a direct target of our series.

**Figure 2.**
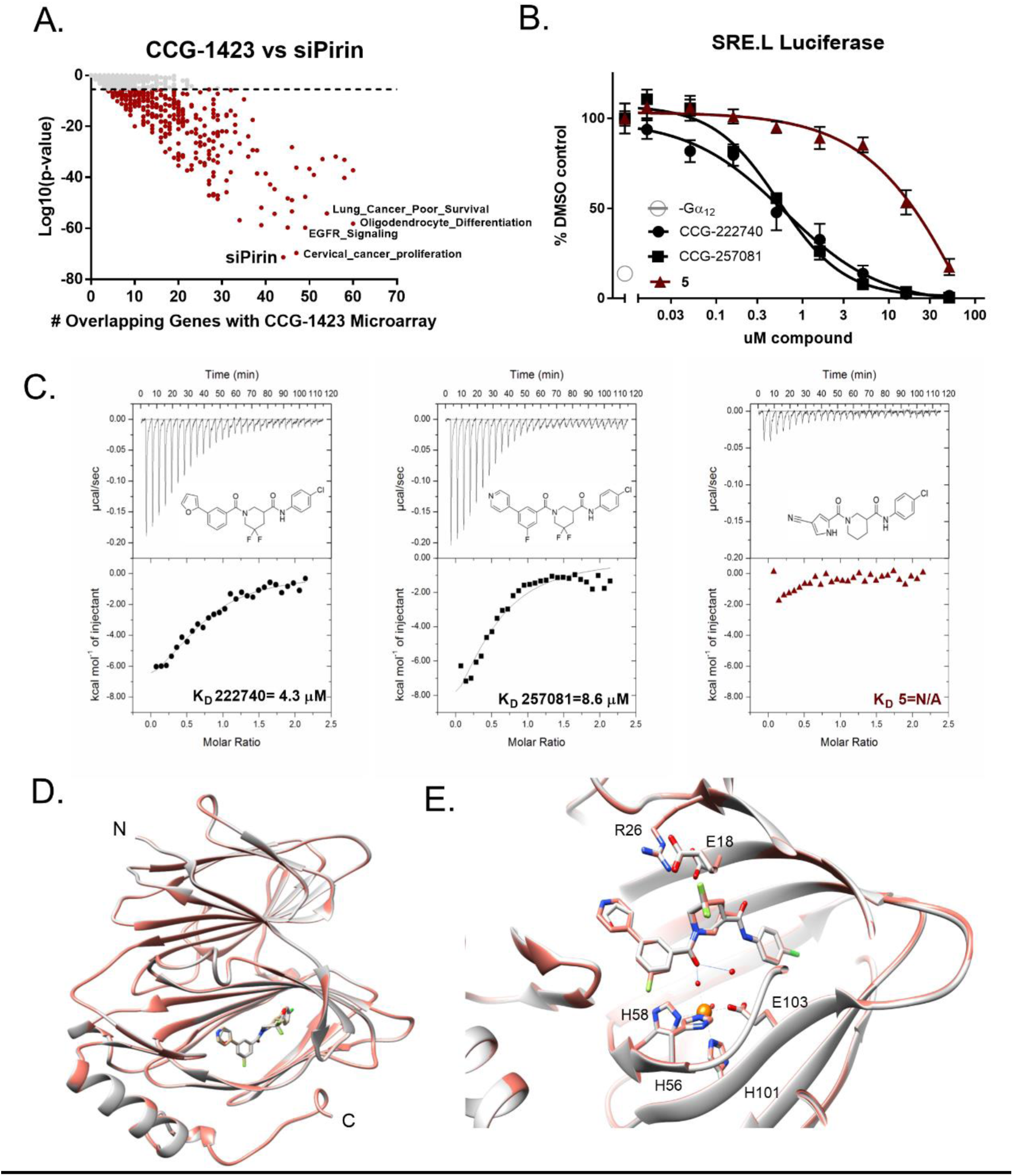
Pirin is a molecular target of CCG-222740 and CCG-257081. (A)Differential gene expression (based on fold change) was calculated for Pirin knockdown (GSE 17551) and CCG-1423 treatment (GSE 30188) and found to have high significance and an overlap of 44 genes in their datasets. (B)HEK293T cells co-transfected with Q231L Gα_12_ and a SRE.L Luciferase reporter were treated with CCG-222740, CCG-257081 or CCG-258531 (C) Isotherm generated by ITC shows that CCG-257081 (black) has a greater enthalpy change upon binding to pirin, as compared to CCG-258531 (red), indicative of better binding to recombinant pirin. (D) Crystal structure overlap of pirin bound to CCG-222740 (gray) and CCG-257081 (salmon). (E) Detailed view of the compound binding pocket with overlaid CCG-222740 (gray) and CCG-257081 (salmon) pirin-bound structures. Also indicated is the metal ion (orange) as well as coordinating residues and water molecules (dashed lines), and hydrogen bonds (solid lines).

To validate pirin as a molecular target, recombinant pirin was purified in order to test whether CCG-222740 and CCG-257081 engaged with pirin *in vitro*. As a comparison, we also tested **5**, a structurally related inhibitor that was nearly inactive in our Gα_12_ induced SRE.L Luciferase assay (Figure 2B and C). Purified, recombinant pirin was used in isothermal titration calorimetery (ITC) experiments to determine the K_D_ of our compounds. CCG-222740 and CCG-257081 bound to pirin with a K_D_ of 4.3 μM and 8.5 μM, respectively. The change in enthalpy for the ‘inactive’ compound **5** was much less than that seen for either CCG-222740 or CCG-257081; it did not provide a reliable fit for binding analysis. Although this is a small set of compounds, it is encouraging to note that an inactive compound in our cell-based assay shows minimal binding to pirin *in vitro* (Figure 2C).

Furthermore, to understand how our small molecules bind to pirin, we solved high-resolution x-ray co-crystal structures of pirin in complex with CCG-222740 and CCG-257081 (1.7 Å and 1.5 Å, respectively) (Figure 2D). The two compounds bind very similarly, with the 4-chloroaniline projecting deep into the hydrophobic pocket and the furan/pyridine rings projecting out into solvent (Figure 2D and Figure 2E). There are no direct contacts between the compound and the iron; the interaction is mostly mediated by hydrogen bonds with Feligated waters (Figure 2E). When these compound-bound pirin co-crystal structures are compared to both the inactive (Fe^2+^ bound), and the active (Fe^3+^ bound) forms described in the literature, a high level of similarity exists. There is an overall RMSD between 257081-bound pirin and Fe^2+^ bound pirin of 0.251 Å and between 257081-bound pirin and Fe^3+^ bound pirin of 0.582 Å. Overall, these results support the idea that pirin is a molecular target of the CCG-222740 series of compounds.

### Modulation of pirin can affect MRTF/SRF dependent SRE.L Luciferase

To explore the connection between pirin and MRTF/SRF signaling, we tested whether the literature pirin inhibitor, CCT251236, could inhibit Gα_12_ mediated SRE.L Luciferase, the same assay used to discover CCG-1423. The SRE.L Luciferase reporter contains several SRF binding sites that were modified to be dependent on MRTF, but lack responsiveness to ETS factors.^1^ CCT251236 has nM potency in a cell-based assay to identify inhibitors of Heat Shock Factor 1 transcription and binds to pirin *in vitro* with a K_D_ of 44 nM as measured by SPR.^28^ CCT251236 had a remarkable effect on SRE.L luciferase expression, with an IC_50_ of 3.3 nM, without affecting luciferase catalytic activity directly (Figure 3A and Supplemental Figure 3). This pharmacologically validates pirin in the same Gα_12_ driven SRE.L Luciferase assay that we discovered and developed our series of compounds. To determine whether overexpression of pirin alone can modulate SRE.L dependent luciferase, pirin was overexpressed in HEK293T cells with a SRE.L Luciferase reporter. Pirin alone was capable of modestly increasing SRE.L driven luciferase expression, suggesting that pirin can enhance MRTF/SRF transcription (Figure 3B). Moreover, to determine whether reduction of pirin through siRNA could suppress MRTF-A-dependent SRE.L Luciferase, primary human dermal fibroblasts were treated with a Dharmacon smartpool siRNA targeting pirin. MRTF-A-driven SRE.L Luciferase was significantly reduced when pirin mRNA levels were decreased. This suggests that pirin modulates gene transcription at the MRTF-A level of regulation (Figure 3C and D). Taken together, these results support an interplay between pirin and MRTF-and SRF-driven luciferase transcription.

**Figure 3.**
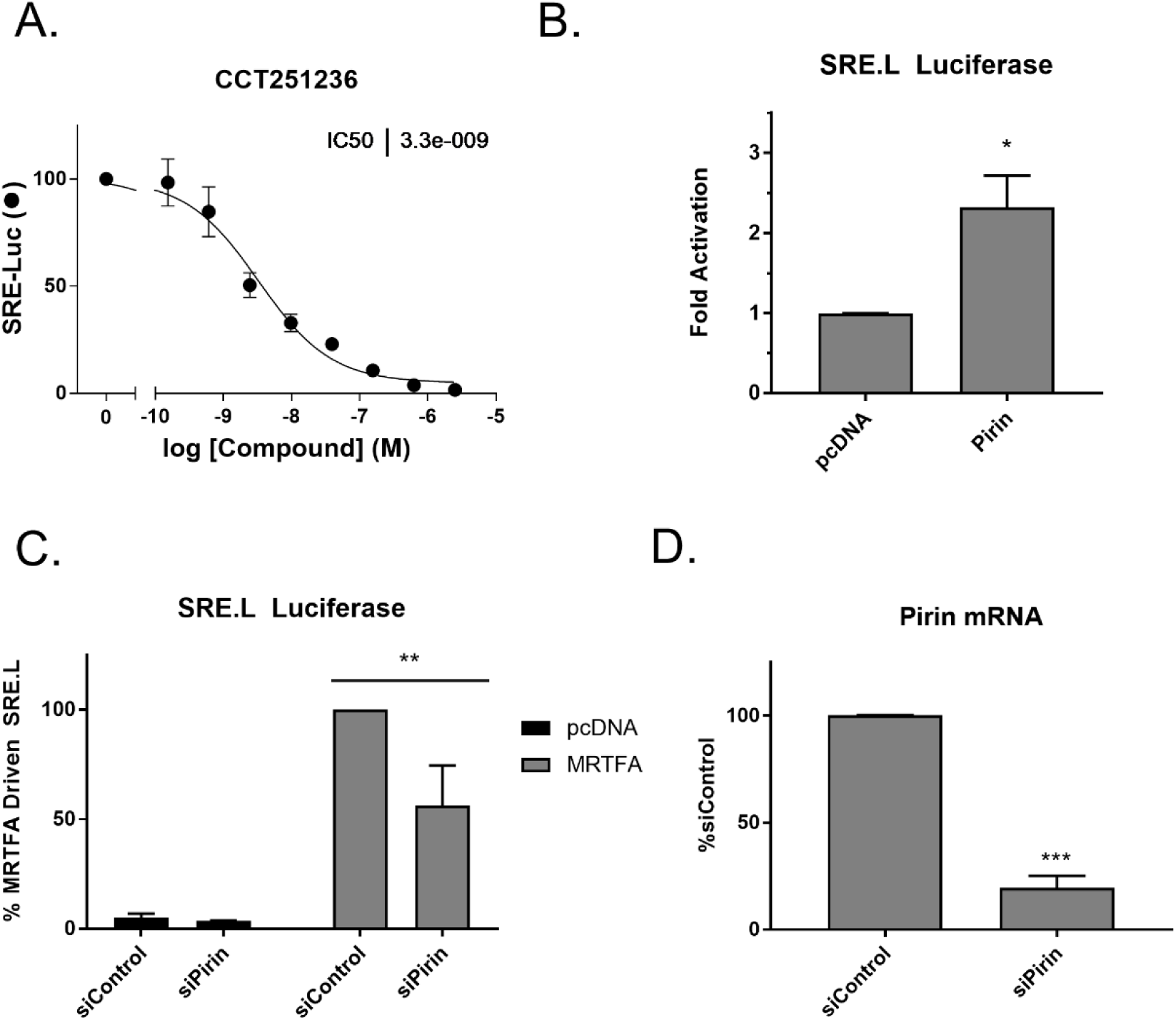
Pirin interacts with the MRTF/SRF pathway. (A) CCT251236 inhibits Q231L Gα_12_ activated SRE.L Luciferase in HEK293T cells;n=4 (B) Overexpression of C-terminally V5 tagged pirin induces SRE.L Luciferase signal in HEK293T cells. Results are expressed as the mean ± SEM. ***p< 0.05 using Student’s t-test; n=3 (C) Knockdown of pirin also reduces MRTF-A driven SRE.L Luciferase in primary dermal fibroblasts. Results are expressed as the mean ±SEM **p<0.01 using 2-way ANOVA; n=3 (D) Validation of pirin knockdown using qPCR. Results are expressed as the mean ± SEM ***p<0.001 using Student’s t-test; n=3.

### Inhibition or ablation of pirin reduces TGF-B induced profibrotic gene expression

The CCG-203971/CCG-222740 series is known to inhibit pro-fibrotic signaling and a large body of literature exists that suggests this is through MRTF/SRF.^13,16,30–31^ However, to pharmacologically verify that pirin is involved in TGF-β dependent gene expression, we used CCT251236, the previously published pirin inhibitor.^28^ Human primary dermal fibroblasts were activated with TGF-β for 24 hours with or without CCG-222740 or CCG-257081 as well as CCT251236. All three compounds significantly reduced TGFβ induced ACTA2 expression (Figure 4A). ACTA2 is the gene for a-Smooth Muscle Actin (α-SMA), which is a marker of myofibroblast transition, and a direct target of MRTF/SRF-mediated transcription^31^. In addition, we tested these compounds against CTGF expression (CCN2). CTGF is a pro-fibrotic mediator that is also, in part, regulated by MRTF/SRF.^5^ We also observed a decrease in CTGF mRNA levels after compound treatment (Figure 4B).

**Figure 4:**
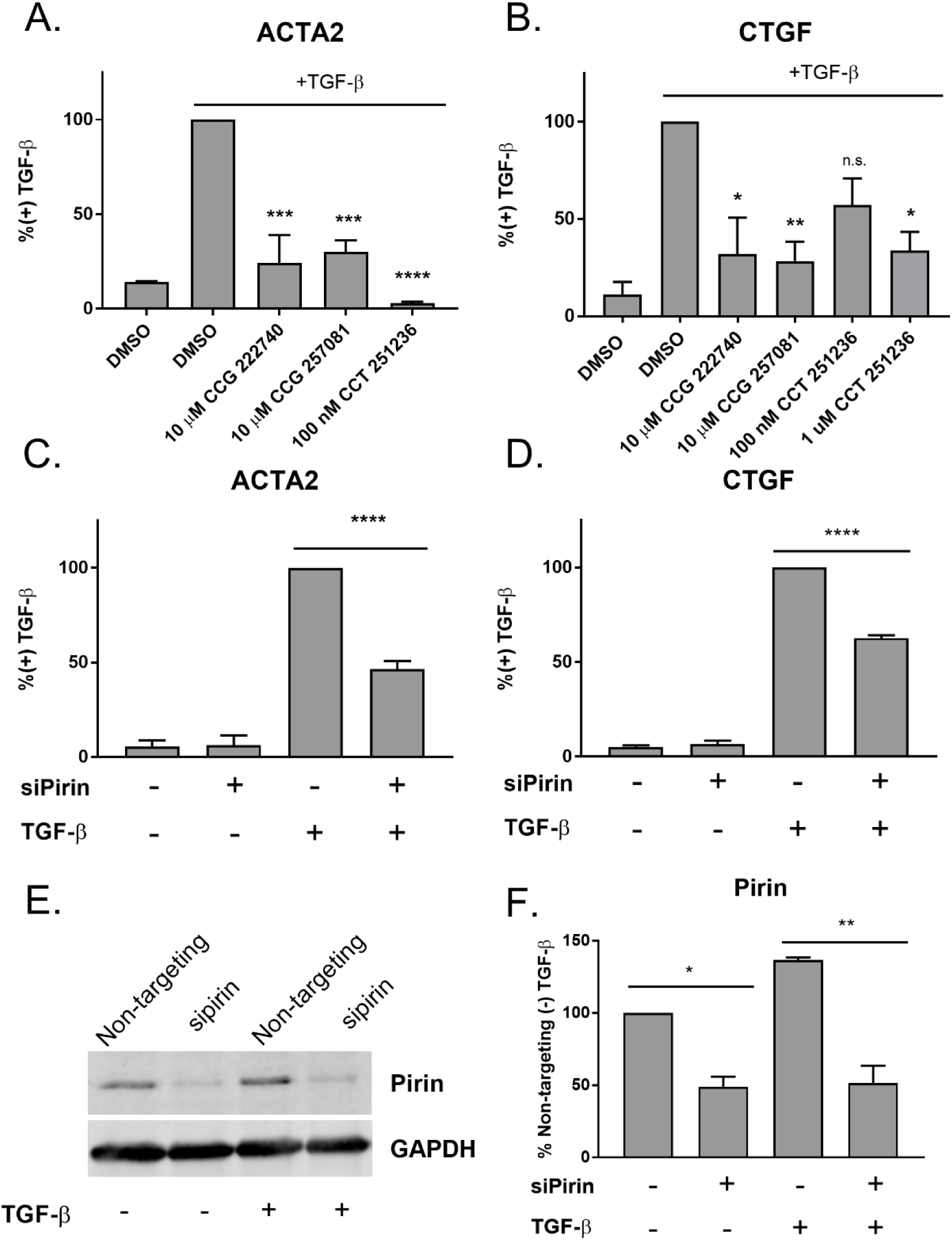
Inhibition or ablation of pirin reduces TGF-β dependent gene expression. (A) Primary dermal fibroblasts from healthy donors were treated with TGF-β and either vehicle control, CCG compounds, or CCT251236. Levels of ACTA2 were measured by qPCR. Results are expressed as the mean ± SEM. ***p< 0.001, ****p<0.0001 using One-way ANOVA; n=3 (B). Similarly, CTGF mRNA levels were measured after treatment with TGF-β and pirin inhibitors. Results are expressed as the mean ± SEM. *p<0.05, **p<0.01 using One-way ANOVA; n=3. (C) Pirin knockdown reduces TGF-β stimulated ACTA2 in human primary dermal fibroblasts. Results are expressed as the mean ± SEM ****p<0.0001 using One-way ANOVA; n=3. Non-targeting siRNA was used in siPirin (-) conditions. (D) Pirin knockdown reduces TGF-β stimulated CTGF. Primary dermal fibroblasts were treated similarly to C. The results are expressed as the mean ± SEM of ****p=0.0001; n=3. (E) Western blot of pirin protein after siRNA treatment. The results are expressed as the mean ± SEM of two independent experiments (*p<0.05, **p<0.01, using One-way ANOVA). (F) Quantification of E.

To further confirm that pirin is involved in TGF-β mediated gene expression, we reduced pirin expression through siRNA. Knockdown of pirin led to a 50% decrease in TGF-β induced ACTA2 and CTGF mRNA, further verifying the role of pirin in TGF-β mediated gene expression (Figure 4C and 4D). Pirin mRNA levels were reduced 80% in primary dermal fibroblasts after siRNA treatment (Supplemental Figure 4) and protein amounts were reduced approximately 50% (Figure 4E and 4F)

### CCG-257081 prevents bleomycin induced skin fibrosis

We have previously shown that CCG-203971^13^ and CCG-232601^17^ can prevent bleomycin-induced fibrosis in mice. However, the most recently developed compound in this series, CCG-257081, which has improved pharmacokinetics, has not yet been tested *in vivo*.^17^ Therefore, CCG-257081 was tested in a bleomycin-induced skin prevention model in mice at 50 mg/kg/day. Fibrosis was induced with daily bleomycin injections and mice were treated with CCG-257081 (50 mg/kg/day by oral gavage) for 14 days. At the conclusion of the experiment, mice were sacrificed and the skin was paraffin-embedded. Tissue slices were used for histological analysis by Masson’s trichrome staining. As shown in Figure 5, treatment of mice with CCG-257081 significantly reduced skin thickening and total collagen content (Figure 5).

**Figure 5.**
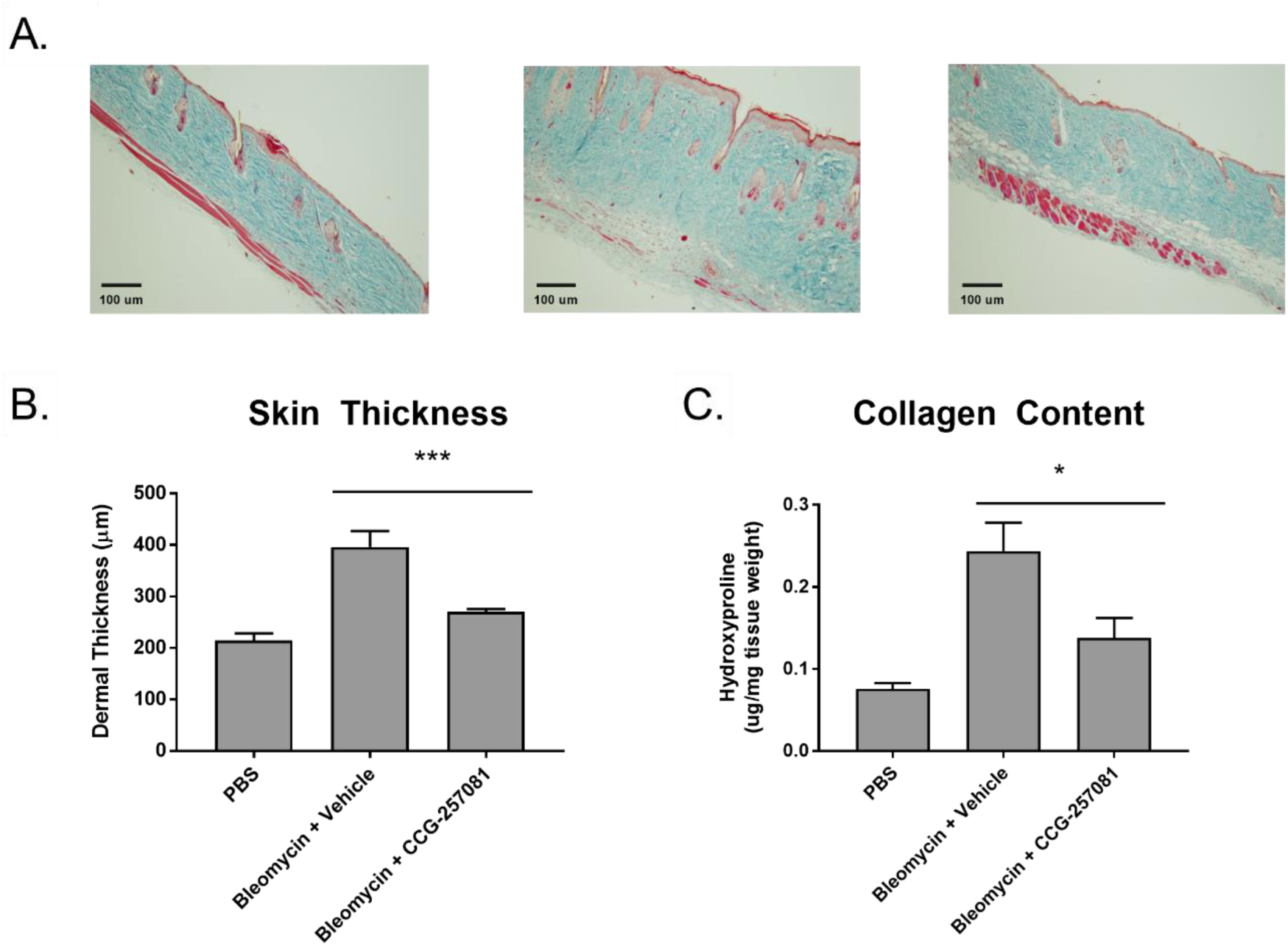
CCG-257081 prevents bleomycin induced fibrosis. Mice treated with 50 mg/kg CCG-257081 had significantly reduced skin thickness as compared to vehicle control. (A) Masson’s trichrome stained skin sections from vehicle treated mice (left panel) bleomycin treated mice (middle panel) and bleomycin+CCG-257081 treated mice (right panel). N=7 (B) Quantification of (A) by measuring the maximal distance between the epidermal-dermal junction and the dermal-subcutaneous fat junction ***p<0.001 using One-way ANOVA. (C) Quantification of collagen using hydroxyproline measurements *p<0.05 using One-way ANOVA.

The CCG-series of compounds has been published in many papers and has demonstrated efficacy in a wide range of disease models. Although these compounds were originally identified in a cell-based luciferase assay that relied on MRTF/SRF-regulated gene transcription, no molecular target was fully validated. Our own preliminary work failed to confirm the report that our series of compounds could inhibit an actin:MRTF interaction^18^ (Supplemental Figure 5), suggesting that the RPEL domain of MRTF was not a direct target of these compounds. In this study, we describe the identification and validation of pirin as a molecular target of CCG-222740 and CCG-257081.

Further work will yield insights into the biology of pirin and how it relates to the MRTF/SRF signaling pathway and pro-fibrotic mechanisms. Pirin has been implicated in melanoma migration and progression^25^, ^32–33^ but has not been explored in terms of fibrotic disease. This is the first report implicating pirin in a pro-fibrotic signaling pathway, so dissecting the exact role of pirin in TGF-β mechanisms will be an important question to address in the future.

The exact mechanism of compound effects on pirin activity is also unknown. It is possible that these compounds stabilize either an oxidized or reduced form of pirin, and therefore affect the ability of pirin to bind to co-activators or transcription factors; it has been shown that only Fe^3+^ bound pirin will bind to p65, whereas Fe^2+^ bound will no.t^26^ Also, CCG-1423 has been shown to displace MRTF-A and SRF from the ACTA2 promoter^34^ which suggests that compound binding to pirin may disrupt a pirin/MRTF-A/SRF/DNA complex. Future work will address these and other questions, including exploring the implications of pirin modulation on global measures of gene transcription.

## Materials and Methods

For Materials and Methods, see Supporting Information.

## Acknowledgements

This research was supported by NIH NIAMS award R01AR066049 (SDL) and NIH NIGMS award R01GM115459-01A1 (RRN). IG was supported by the Spartan Innovation Fund. This research used resources of the Advanced Photon Source, a U.S. Department of Energy (DOE) Office of Science User Facility operated for the DOE Office of Science by Argonne National Laboratory under Contract No. DE-AC02-06CH11357. Use of the LS-CAT Sector 21 was supported by the Michigan Economic Development Corporation and the Michigan Technology Tri-Corridor (Grant 085P1000817). Molecular graphics and analyses were performed with the UCSF Chimera package. Chimera is developed by the Resource for Biocomputing, Visualization, and Informatics at the University of California, San Francisco (supported by NIGMS P41-GM103311).

## References

1. Hill, C. S.; Wynne, J.; Treisman, R., The Rho family GTPases RhoA, Rac1, and CDC42Hs regulate transcriptional activation by SRF. Cell. 1995, 81 (7), 1159–70.

2. Olson, E. N.; Nordheim, A., Linking actin dynamics and gene transcription to drive cellular motile functions. Nat Rev Mol Cell Biol. 2010, 11 (5), 353–65.

3. Gualdrini, F.; Esnault, C.; Horswell, S.; Stewart, A.; Matthews, N.; Treisman, R., SRF Co-factors Control the Balance between Cell Proliferation and Contractility. Mol Cell. 2016, 64 (6), 1048–1061. doi:10.1016/j.molcel.2016.10.016. Epub 2016 Nov 17.

4. Tsou, P. S.; Haak, A. J.; Khanna, D.; Neubig, R. R., Cellular mechanisms of tissue fibrosis. 8. Current and future drug targets in fibrosis: focus on Rho GTPase-regulated gene transcription. Am J Physiol Cell Physiol. 2014, 307 (1), C2–13. doi:10.1152/ajpcell.00060.2014. Epub 2014 Apr 16.

5. Medjkane, S.; Perez-Sanchez, C.; Gaggioli, C.; Sahai, E.; Treisman, R., Myocardin-related transcription factors and SRF are required for cytoskeletal dynamics and experimental metastasis. Nat Cell Biol 2009, 11 (3), 257–68.

6. Evelyn, C. R.; Wade, S. M.; Wang, Q.; Wu, M.; Iniguez-Lluhi, J. A.; Merajver, S. D.; Neubig, R. R., CCG-1423: a small-molecule inhibitor of RhoA transcriptional signaling. Mol Cancer Ther 2007, 6 (8), 2249–60.

7. Gau, D.; Veon, W.; Capasso, T. L.; Bottcher, R.; Shroff, S.; Roman, B. L.; Roy, P., Pharmacological intervention of MKL/SRF signaling by CCG-1423 impedes endothelial cell migration and angiogenesis. Angiogenesis. 2017, 20 (4), 663–672. doi:10.1007/s10456-017-9560-y. Epub 2017 Jun 21.

8. Jin, W.; Goldfine, A. B.; Boes, T.; Henry, R. R.; Ciaraldi, T. P.; Kim, E. Y.; Emecan, M.; Fitzpatrick, C.; Sen, A.; Shah, A.; Mun, E.; Vokes, V.; Schroeder, J.; Tatro, E.; Jimenez-Chillaron, J.; Patti, M. E., Increased SRF transcriptional activity in human and mouse skeletal muscle is a signature of insulin resistance. J Clin Invest. 2011, 121 (3), 918–29. doi:10.1172/JCI41940.

9. Mae, S.; Shirasawa, S.; Yoshie, S.; Sato, F.; Kanoh, Y.; Ichikawa, H.; Yokoyama, T.; Yue, F.; Tomotsune, D.; Sasaki, K., Combination of small molecules enhances differentiation of mouse embryonic stem cells into intermediate mesoderm through BMP7-positive cells. Biochem Biophys Res Commun. 2010, 393 (4), 877–82. doi:10.1016/j.bbrc.2010.02.111. Epub 2010 Feb 19.

10. Sakai, N.; Chun, J.; Duffield, J. S.; Wada, T.; Luster, A. D.; Tager, A. M., LPA1-induced cytoskeleton reorganization drives fibrosis through CTGF-dependent fibroblast proliferation. FASEB J. 2013, 27 (5), 1830–46. doi:10.1096/fj.12-219378. Epub 2013 Jan 15.

11. Evelyn, C. R.; Bell, J. L.; Ryu, J. G.; Wade, S. M.; Kocab, A.; Harzdorf, N. L.; Hollis Showalter, H. D.; Neubig, R. R.; Larsen, S. D., Design, synthesis and prostate cancer cell-based studies of analogs of the Rho/MKL1 transcriptional pathway inhibitor, CCG-1423. Bioorg Med Chem Lett 2010, 20 (2), 665–72.

12. Bell, J. L.; Haak, A. J.; Wade, S. M.; Kirchhoff, P. D.; Neubig, R. R.; Larsen, S. D., Optimization of novel nipecotic bis(amide) inhibitors of the Rho/MKL1/SRF transcriptional pathway as potential anti-metastasis agents. Bioorg Med Chem Lett. 2013, 2S (13), 3826–32. doi:10.1016/j.bmcl.2013.04.080. Epub 2013 May 7.

13. Haak, A. J.; Tsou, P. S.; Amin, M. A.; Ruth, J. H.; Campbell, P.; Fox, D. A.; Khanna, D.; Larsen, S. D.; Neubig, R. R., Targeting the myofibroblast genetic switch: inhibitors of myocardin-related transcription factor/serum response factor-regulated gene transcription prevent fibrosis in a murine model of skin injury. J Pharmacol Exp Ther. 2014, S49 (3), 480–6. doi:10.1124/jpet. 114.213520. Epub 2014 Apr 4.

14. Sisson, T. H.; Ajayi, I. O.; Subbotina, N.; Dodi, A. E.; Rodansky, E. S.; Chibucos, L. N.; Kim, K. K.; Keshamouni, V. G.; White, E. S.; Zhou, Y.; Higgins, P. D.; Larsen, S. D.; Neubig, R. R.; Horowitz, J. C., Inhibition of myocardin-related transcription factor/serum response factor signaling decreases lung fibrosis and promotes mesenchymal cell apoptosis. Am J Pathol. 2015, 185 (4), 969–86. doi:10.1016/j.ajpath.2014.12.005. Epub 2015 Feb 11.

15. Haak, A. J.; Appleton, K. M.; Lisabeth, E. M.; Misek, S. A.; Ji, Y.; Wade, S. M.; Bell, J. L.; Rockwell, C. E.; Airik, M.; Krook, M. A.; Larsen, S. D.; Verhaegen, M.; Lawlor, E. R.; Neubig, R. R., Pharmacological Inhibition of Myocardin-related Transcription Factor Pathway Blocks Lung Metastases of RhoC-Overexpressing Melanoma. Mol Cancer Ther. 2017, 16 (1), 193–204. doi:10.1158/1535-7163.MCT-16-0482. Epub 2016 Nov 11.

16. Yu-Wai-Man, C.; Spencer-Dene, B.; Lee, R. M.; Hutchings, K.; Lisabeth, E. M.; Treisman, R.; Bailly, M.; Larsen, S. D.; Neubig, R. R.; Khaw, P. T., Local delivery of novel MRTF/SRF inhibitors prevents scar tissue formation in a preclinical model of fibrosis. Sci Rep. 2017, 7 (1), 518. doi:10.1038/s41598-017-00212-w.

17. Hutchings, K. M.; Lisabeth, E. M.; Rajeswaran, W.; Wilson, M. W.; Sorenson, R. J.; Campbell, P. L.; Ruth, J. H.; Amin, A.; Tsou, P. S.; Leipprandt, J. R.; Olson, S. R.; Wen, B.; Zhao, T.; Sun, D.; Khanna, D.; Fox, D. A.; Neubig, R. R.; Larsen, S. D., Pharmacokinetic optimitzation of CCG-203971: Novel inhibitors of the Rho/MRTF/SRF transcriptional pathway as potential antifibrotic therapeutics for systemic scleroderma. Bioorg Med Chem Lett. 2017, 27 (8), 1744–1749. doi:10.1016/j.bmcl.2017.02.070. Epub 2017 Mar 10.

18. Hayashi, K.; Watanabe, B.; Nakagawa, Y.; Minami, S.; Morita, T., RPEL proteins are the molecular targets for CCG-1423, an inhibitor of Rho signaling. PLoS One. 2014, 9 (2), e89016. doi:10.1371/journal.pone.0089016. eCollection 2014.

19. Lundquist, M. R.; Storaska, A. J.; Liu, T. C.; Larsen, S. D.; Evans, T.; Neubig, R. R.; Jaffrey, S. R., Redox modification of nuclear actin by MICAL-2 regulates SRF signaling. Cell. 2014, 156 (3), 563–76. doi:10.1016/j.cell.2013.12.035. Epub 2014 Jan 16.

20. Bell, J. L.; Haak, A. J.; Wade, S. M.; Sun, Y.; Neubig, R. R.; Larsen, S. D., Design and synthesis of tag-free photoprobes for the identification of the molecular target for CCG-1423, a novel inhibitor of the Rho/MKL1/SRF signaling pathway. Beilstein J Org Chem. 2013, 9:966–7S. (doi), 10.3762/bjoc.9.111. Print 2013.

21. Evelyn, C. R.; Lisabeth, E. M.; Wade, S. M.; Haak, A. J.; Johnson, C. N.; Lawlor, E. R.; Neubig, R. R., Small-Molecule Inhibition of Rho/MKL/SRF Transcription in Prostate Cancer Cells: Modulation of Cell Cycle, ER Stress, and Metastasis Gene Networks. Microarrays (Basel). 2016, 5(2). (pii), E13. doi:10.3390/microarrays5020013.

22. Davda, D.; El Azzouny, M. A.; Tom, C. T.; Hernandez, J. L.; Majmudar, J. D.; Kennedy, R. T.; Martin, B. R., Profiling targets of the irreversible palmitoylation inhibitor 2-bromopalmitate. ACS Chem Biol. 2013, 8 (9), 1912–7. doi:10.1021/cb400380s. Epub 2013 Jul 25.

23. Wendler, W. M.; Kremmer, E.; Forster, R.; Winnacker, E. L., Identification of pirin, a novel highly conserved nuclear protein. J Biol Chem. 1997, 272 (13), 8482–9.

24. Dechend, R.; Hirano, F.; Lehmann, K.; Heissmeyer, V.; Ansieau, S.; Wulczyn, F. G.; Scheidereit, C.; Leutz, A., The Bcl-3 oncoprotein acts as a bridging factor between NF-kappaB/Rel and nuclear co-regulators. Oncogene. 1999, 18 (22), 3316–23. doi:10.1038/sj.onc.1202717.

25. Miyazaki, I.; Simizu, S.; Okumura, H.; Takagi, S.; Osada, H., A small-molecule inhibitor shows that pirin regulates migration of melanoma cells. Nat Chem Biol. 2010, 6 (9), 667–73. doi:10.1038/nchembio.423. Epub 2010 Aug 15.

26. Liu, F.; Rehmani, I.; Esaki, S.; Fu, R.; Chen, L.; de Serrano, V.; Liu, A., Pirin is an iron-dependent redox regulator of NF-kappaB. Proc Natl Acad Sci U S A. 2013, 11G (24), 9722–7. doi:10.1073/pnas. 1221743110. Epub 2013 May 28.

27. Brzoska, K.; Stepkowski, T. M.; Kruszewski, M., Basal PIR expression in HeLa cells is driven by NRF2 via evolutionary conserved antioxidant response element. Mol Cell Biochem. 2014, SB9 (1-2), 99–111. doi:10.1007/s11010-013-1931-0. Epub 2014 Jan 5.

28. Cheeseman, M. D.; Chessum, N. E.; Rye, C. S.; Pasqua, A. E.; Tucker, M. J.; Wilding, B.; Evans, L. E.; Lepri, S.; Richards, M.; Sharp, S. Y.; Ali, S.; Rowlands, M.; O’Fee, L.; Miah, A.; Hayes, A.; Henley, A. T.; Powers, M.; Te Poele, R.; De Billy, E.; Pellegrino, L.; Raynaud, F.; Burke, R.; van Montfort, R. L.; Eccles, S. A.; Workman, P.; Jones, K., Discovery of a Chemical Probe Bisamide (CCT251236): An Orally Bioavailable Efficacious Pirin Ligand from a Heat Shock Transcription Factor 1 (HSF1) Phenotypic Screen. J Med Chem. 2017, 6G (1), 180–201. doi:10.1021/acs.jmedchem.6b01055. Epub 2016 Dec 22.

29. Johnson, L. A.; Rodansky, E. S.; Haak, A. J.; Larsen, S. D.; Neubig, R. R.; Higgins, P. D., Novel Rho/MRTF/SRF inhibitors block matrix-stiffness and TGF-beta-induced fibrogenesis in human colonic myofibroblasts. Inflamm Bowel Dis. 2014, 2G (1), 154–65. doi:10.1097/01.MIB.0000437615.98881.31.

30. Crider, B. J.; Risinger, G. M., Jr.; Haaksma, C. J.; Howard, E. W.; Tomasek, J. J., Myocardin-related transcription factors A and B are key regulators of TGF-beta1-induced fibroblast to myofibroblast differentiation. J Invest Dermatol. 2011, 1S1 (12), 2378–85. doi:10.1038/jid.2011.219. Epub 2011 Jul 21.

31. Speight, P.; Kofler, M.; Szaszi, K.; Kapus, A., Context-dependent switch in chemo/mechanotransduction via multilevel crosstalk among cytoskeleton-regulated MRTF and TAZ and TGFbeta-regulated Smad3. Nat Commun. 2016, 7:11642. (doi), 10.1038/ncomms11642.

32. Licciulli, S.; Luise, C.; Scafetta, G.; Capra, M.; Giardina, G.; Nuciforo, P.; Bosari, S.; Viale, G.; Mazzarol, G.; Tonelli, C.; Lanfrancone, L.; Alcalay, M., Pirin inhibits cellular senescence in melanocytic cells. Am J Pathol. 2011, 178 (5), 2397–406. doi:10.1016/j.ajpath.2011.01.019.

33. Licciulli, S.; Luise, C.; Zanardi, A.; Giorgetti, L.; Viale, G.; Lanfrancone, L.; Carbone, R.; Alcalay, M., Pirin delocalization in melanoma progression identified by high content immuno-detection based approaches. BMC Cell Biol. 2010, 11:5. (doi), 10.1186/1471-2121-11-5.

34. Foster, C. T.; Gualdrini, F.; Treisman, R., Mutual dependence of the MRTF-SRF and YAP-TEAD pathways in cancer-associated fibroblasts is indirect and mediated by cytoskeletal dynamics. Genes Dev. 2017, 31 (23-24), 2361-2375. doi:10.1101/gad.304501.117. Epub 2018 Jan 9.

